# Remodeling of the core leads HIV-1 pre-integration complex in the nucleus of human lymphocytes

**DOI:** 10.1101/2020.01.24.918383

**Authors:** Guillermo Blanco-Rodriguez, Anastasia Gazi, Blandine Monel, Stella Frabetti, Viviana Scoca, Florian Mueller, Olivier Schwartz, Jacomine Krijnse-Locker, Pierre Charneau, Francesca Di Nunzio

## Abstract

Retroviral replication proceeds through obligate integration of the viral DNA into the host genome. To enter the nucleus, the viral DNA must be led through the nuclear pore complex (NPC). During HIV-1 cytoplasmic journey, the viral core acts like a shell to protect the viral genetic material from antiviral sensors and ensure an adequate environment for the reverse transcription. However, the relatively narrow size of the nuclear pore channel requires that the HIV-1 core reshapes into a structure that fits the pore. On the other hand, the organization of the viral CA proteins that remain associated to the pre-integration complex (PIC) during and after nuclear translocation, in particular, in human lymphocytes, the main target cells of HIV-1, is still enigmatic. In this study, we analysed the progressive organizational changes of viral CA proteins within the cytoplasm and the nucleus by immuno-gold labelling. Furthermore, we set up a novel technology, HIV-1 ANCHOR, which enables specific detection of the retrotranscribed DNA by fluorescence microscopy, thereby uncovering the architecture of the potential HIV-1 PIC. Thus, we revealed DNA- and CA-positive complexes by correlated light- and electron microscopy (CLEM). During and after nuclear translocation, HIV-1 appears as a complex of viral DNA decorated by multiple viral CA proteins remodelled in a “pearl necklace” shape. Thus, we observed how CA proteins reshape around the viral DNA to permit the entrance of the HIV-1 in the nucleus. This particular CA protein complex composed by the integrase and the retrotranscribed DNA leads HIV-1 genome inside the host nucleus to potentially replicate.

Our findings contribute to the understanding of the early steps of HIV-1 infection and provide new insights into the organization of HIV-1 CA proteins during and after viral nuclear entry.

**Importance:** How the reverse transcribed genome reaches the host nucleus remains a main open question related to the infectious cycle of HIV-1. HIV-1 core has a size of ∼100 nm, largely exceeding that of the NPC channel (∼39 nm). Thus, a rearrangement of the viral CA proteins organization is required to achieve effective nuclear translocation. The mechanistic of this process remains undefined due to the lack of a technology capable to visualize potential CA sub-complexes in association with the viral DNA in the nucleus of HIV-1-infected cells.

By the means of state-of-the-art technologies (HIV-1 ANCHOR system combined with CLEM), our study shows that remodeled viral complexes retain multiple CA proteins but not intact core or only a single CA monomer. These viral CA complexes associated with the retrotranscribed DNA can be observed in the outer and inner side of the NE, and they represent potential PIC.

Thus, our study shed light on critical early steps characterizing HIV-1 infection, thereby revealing novel, therapeutically exploitable points of intervention. Furthermore, we developed and provided a powerful tool enabling direct, specific and high-resolution visualization of intracellular and intranuclear HIV-1 subviral structures.

## Introduction

Upon viral entry, the HIV-1 core is released into the host cell cytoplasm and starts its journey towards the nucleus (1, 2) helped or hampered by cellular factors (3–10). The viral core has a conical shape (120nm × 60nm × 40 nm), composed by [1500 capsid (CA) monomers organized in hexamers or pentamers (11–13). This conical structure acts as a protective shell for the viral DNA against defence mechanisms of the host, like nucleic acid sensors and nucleases (14–16) and can be the target of restriction factors, such as Trim5α and MX2 (9, 10). Furthermore, the viral CA seems to participate in two critical steps of HIV-1 life cycle, nuclear translocation (4–7, 17) and integration (18–23). The viral core is considered a fragile structure (24) that could exist only as intact or completely unbundled. However, a recent *in vitro* study highlighted the possibility that partial cores can be stabilized by host factors (25). Thus, HIV-1 CA can exist in different forms in infected cells: as intact cores, as monomers and probably as partial cores. Of note, the viral core largely exceeds the size (11–13) of the nuclear pore channel (26); thus, the core should disassemble before entering the nucleus. The “immediate uncoating” model, which postulates a complete loss of the viral CA proteins (2), has been the most accredited model in the past, supported by the impossibility to co-purify viral CA with other viral proteins at early time points due to its genetic fragility (24, 27). Contrary to this model, Aiken’s group (20) was the first to propose a role for CA in post-nuclear entry steps, followed by other studies (21–23, 28). However, the viral CA protein was biochemically detected for the first time in the nucleus of macrophages and HeLa cells by Fassati and colleagues (23). The presence of CA in the nucleus was shown also by the analysis of individual time points of infection in fixed cells using surrogate viruses, or by indirect viral labelling using live imaging (29–38); nonetheless, these studies failed to reveal the organization of CA during viral nuclear translocation. By demonstrating the importance of the CA to regulate and coordinate different steps of HIV-1 life cycle, these findings prompted the development of small molecules (HIV-1 CA inhibitors), thereby promoting the development of a new class of anti-retroviral drugs (39). It should be noted, however, that to date the presence of HIV-1 CA has never been shown in the nucleus of infected primary CD4^+^ T cells, the main target cells of HIV-1. In addition, the CA-positive structure associated with the reverse transcribed DNA that enters the nucleus, in particular in mitotic cells, remains enigmatic. The details of the morphology of intermediate states of viral replication complexes can be analysed only at the nanometre level, due to their small size (HIV-1 core ∼ 100nm)(11). Previous studies showed the possibility to visualize HIV-1 cores in the cytoplasm by CLEM under particular circumstances, such as in the presence of proteasome inhibition (40). Nevertheless, it has been impossible to visualize the intermediate states of viral replication complexes, as well as the organization of the viral PIC during a “quasi” physiological infection at the nanometer level. The viral CA was shown to remain partially associated to the HIV-1 DNA after nuclear entry by confocal microscopy (30). As CA-associated HIV-1 DNA has sub-diffraction size, details on this complex could be revealed only by high resolution imaging, until now unattainable due to the incompatibility of fluorescent DNA labelling techniques, such as DNA FISH or click chemistry (EdU), with transmission electron microscopy (TEM).

Our study provides new evidence on the organization of viral CA proteins before, during and after nuclear translocation. To visualize intermediate states of viral replication complexes, we combined CLEM with a novel fluorescent approach to directly label HIV-1 DNA, which we defined HIV-1 ANCHOR. The combination of both technologies, high resolution electron and fluorescence microscopy, highlighted the configuration of the CA protein in viral complexes during the nuclear translocation step. We identified nuclear viral complexes including the crucial components of a potentially functional PIC, the integrase (IN) and the retrotranscribed DNA.

We reveal that HIV-1 core remodels in multiple CA proteins, as observed in both HeLa cells and primary lymphocytes. Immediately before nuclear translocation, HIV-1 CA complexes reshape in a “pearl necklace” shape, thus enabling the entry of the viral genome in the host nucleus, the cellular compartment of the viral replication.

## Results

### Viral reverse transcription correlates with HIV-1 CA and IN association

We analysed the dynamics of the HIV-1 CA and IN association, based on their colocalization, and their link with viral retrotranscription. As the viral IN cannot be efficiently labelled using a direct antibody, we infected HeLa cells with HIV-1 containing a small HA tag fused to the C terminus of the IN (HIV-1ΔEnv IN_HA_ / VSV-G) (41). The genetically modified virus efficiently infected HeLa cells as well as primary CD4^+^ T lymphocytes similarly to the virus carrying the wild type IN (Fig. 1A, Fig.3B). This feature made this tagged-virus a functional tool to study replication complexes in relation to the structure of the viral capsid. Such HA-labelled IN enabled us to study the association between CA and IN during viral infection using a “quasi” WT virus. Cells fixed at 6 h post-infection showed ∼ 60-70% of colocalization of viral CA with the viral IN (Fig. 1A, Fig.1C). Next, we investigated the importance of the presence of CA and IN in cytoplasmic complexes within infected cells for reverse transcription by using a capsid-targeting small molecule, PF74. PF74 binds at the interface between the N-terminal domain (NTD) and the C-terminal domain (CTD) (19, 42) of CA, thus inhibiting HIV-1 infection by blocking nuclear entry in a reportedly bimodal dose-dependent mechanism (19, 42–44). It has been shown that reverse transcription (RT) can occur in the presence of low dose of PF 74 (< 10 µM), while it was impeded at higher doses of this drug (42, 44). We challenged HeLa cells with HIV-1 in the presence or not of PF74, using low (1.25µM) and high (10 µM) doses. Then, we measured IN/CA colocalization spots per cell at 24h post-infection (Fig. 1B) by immunolabelling the viral CA and IN, respectively. When we applied a low dose of PF74, at 24 h post-infection we could still detect some CA/IN colocalization, this was not the case in samples treated with high dose of PF74 (Fig.1B). In agreement with other studies (19, 44, 45), low dose of PF74 allowed for the reverse transcription to occur (∼60% with respect to the infected sample untreated with PF74) (Fig. 1B). However, since a low dose of PF74 interferes with the nuclear import of HIV-1, viral replication was not detected in these cells, as reported also by others (Fig. 1B) (19, 44, 45). Instead, control cells (untreated with PF74) allow active viral transcription (Fig. 1B). A high dose of PF74 caused a loss of the association between CA and IN, together with a loss of reverse transcription activity (Fig. 1B). We corroborated these results by carrying out dose-response experiments to check for the presence of CA/IN cytoplasmic complexes (6 h post-infection) and viral infectivity by beta-gal assay (48h post-infection) (Fig. 1C). These experiments show a progressive loss of viral CA and viral IN association in a dose-response manner to the PF74 drug. Our data indicate that CA/IN association correlates with viral reverse transcription. (Fig. 1B and C).

**FIG. 1.**
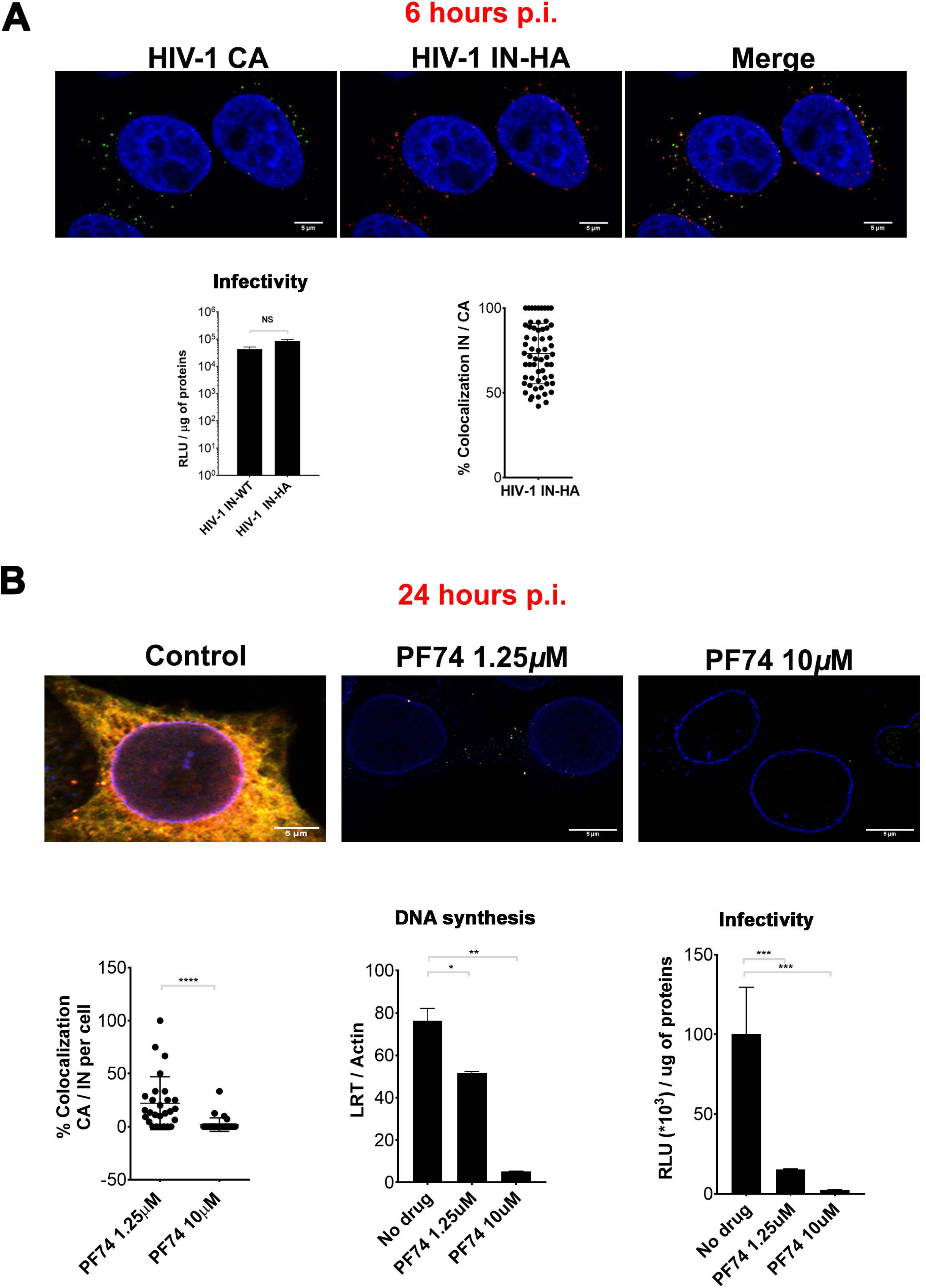
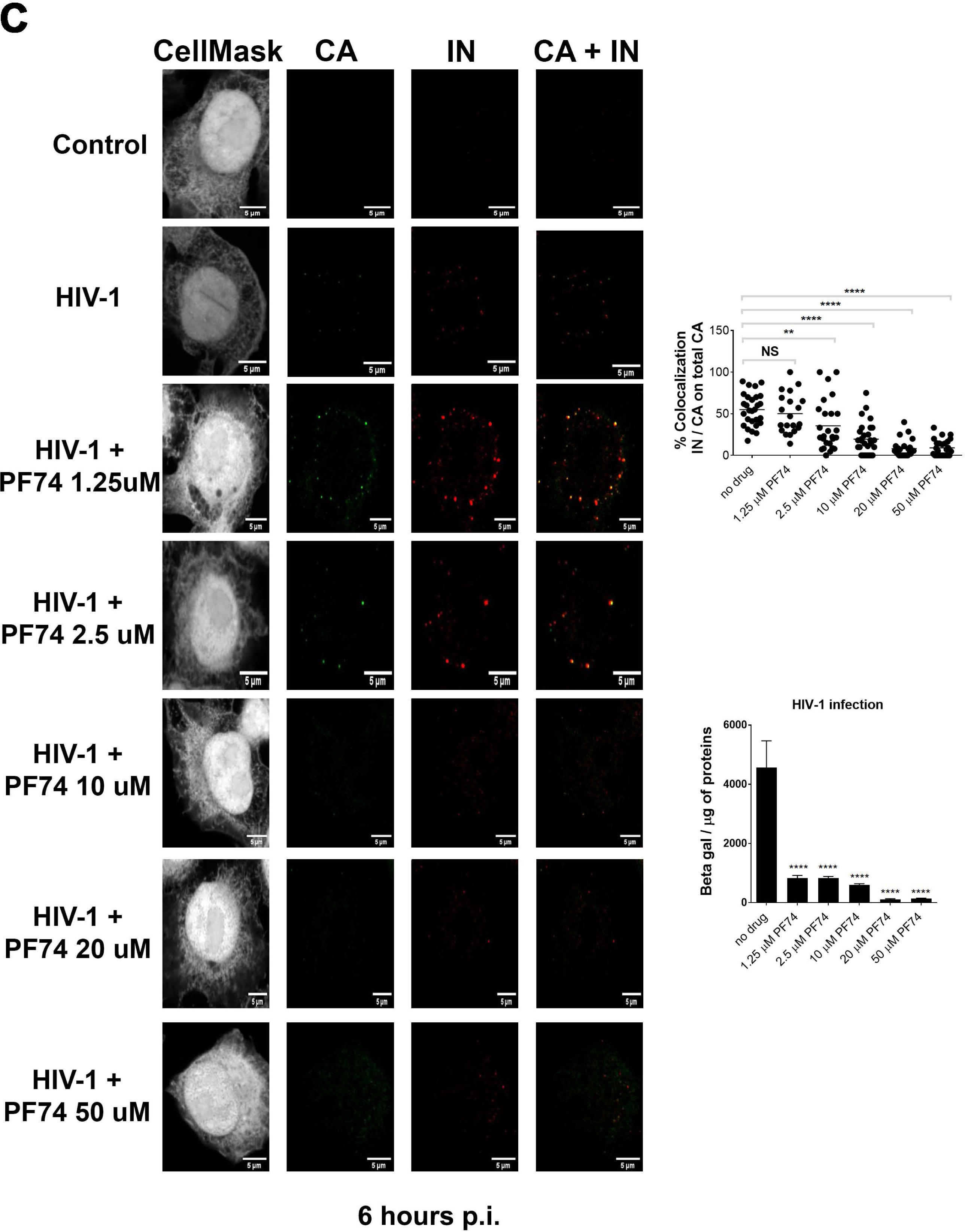
Viral reverse transcription correlates with HIV-1 CA and IN association. (A) HeLa cells (10^6^ cells) infected with 500ng of p24 of HIV-1 Env IN_HA_ / VSV-G fixed at 6 h post infection and labelled with antibodies anti-p24 (green) and anti-HA (red). Analysis of the percentage of IN/CA co-localization analysed by Image J and by Graph Pad Prism 7. Comparison of the infectivity of HIV-1 carrying the IN wild type or the IN fused to HA tag analysed by beta-galactosidase assay, normalized by amount of proteins. (B) HeLa cells infected for 24 h in presence or not of the drug PF74 at low dose (1.25 μ) or high dose (10 μM). Cells were fixed on 4% of PFA and labelled with antibodies anti p24, anti-HA and anti-Nup153. Co-localization between CA and IN was analysed by ImageJ and by Graph Pad Prism 7. DNA synthesis has been evaluated by qPCR through the amplification of late reverse transcripts (LRT). The infectivity was analysed at 48 h post infection by beta galactosidase assay and normalized by protein amount. (C) Effect of PF74 doses on HIV-1 CA and IN detections at 6 h post infection in HeLaP4R5 cells by confocal microscopy. Analysis of the percentage of IN/CA co-localization by Image J and by Graph Pad Prism 7 (graph on the top right). The level of infectivity has been evaluated at 48h post infection by beta galactosidase assay normalized for the quantity of proteins as reported in the histogram on the bottom right. Differences were considered statistically significant at a P value of <0.0001 (****), <0.001 (***), or <0.01 (**), or <0.1 (*), analysed by Graph Pad Prism 7 (*T student test*).

### Remodelling of HIV-1 cores

Next, we sought to visualize in more detail the cytoplasmic structures positive for both CA and IN. We infected HeLa P4R5 cells with 500 ng of p24 per million of cells. The asynchronous HIV-1 infection and the specific time of fixation (6 hours post-infection) allowed visualization of different states of viral replication complexes by EM. We observed several seemingly intact virions residing in endosomes (Fig.2A), consistent with the entry pathway engaged by HIV-1 pseudotyped with VSV-G envelope (46). The intact particles in endosomes contained conical cores, while membrane-free cores, seemingly released from these organelles, were detected in the cytoplasm (Fig. 2A). However, to demonstrate the detection of free cores in the cytoplasm, sections were labelled with anti-CA and a secondary antibody coupled to 10nm gold particles (Fig. 2B). Core like structures that had retained a “quasi” conical shape were also seen in the vicinity of the nuclear envelope (NE). These CA-positive viral structures were usually decorated with 2 gold particles (Fig. 2B-ii,-iii), although the viral core is composed by multiple CA monomers. Uninfected cells labelled with anti-CA and a secondary antibody labelled with gold particles revealed only background labelling, showing the specificity of the antibody (Fig. 2B-i). During and after the nuclear viral entry, conical core like structures were not detected and instead the immuno-labelling decorated a different CA-positive pattern (Fig. 2B). On the cytoplasmic side of the nuclear pore we could detect irregular shapes similar to “core like structures” decorated with anti-CA (Fig. 2B-ii, -iii). In the nucleus, however, the same antibody outlined pearl-necklace shapes, composed by multiple CA proteins (Fig. 2B-iv,-v). Cytoplasmic viral complexes were typically decorated by 2 CA gold particles (∼ 80%) whereas the nuclear complexes were labelled on average by 3 particles (Fig. 2C). The detection of these particular nuclear CA forms was not antibody dependent, because similar results were obtained with another antibody against CA (AG3.0) (Fig. 2D). Importantly, the difference in gold distribution was also seen with wild-type HIV-1 (Fig. 2E), implying that it did not depend on the route of viral entry. By cryo-EM, we confirmed the presence of multiple CA gold particles forming a “pearl necklace” shape inside the nucleus of infected cells (Fig. 2F and G). These elongated CA complexes are better resolved by cryo-EM (47) and detected by more gold particles when a protein A coupled to gold was used instead of a secondary antibody (Fig. 2F). This likely related to a better affinity of the protein A for primary antibodies compared to secondary gold-conjugated antibodies. Importantly, different labelling methods labelled similar structures in the nucleus and confirmed the increase in labelling after nuclear entry (Fig. 2B, D, F and G).

**FIG. 2.**
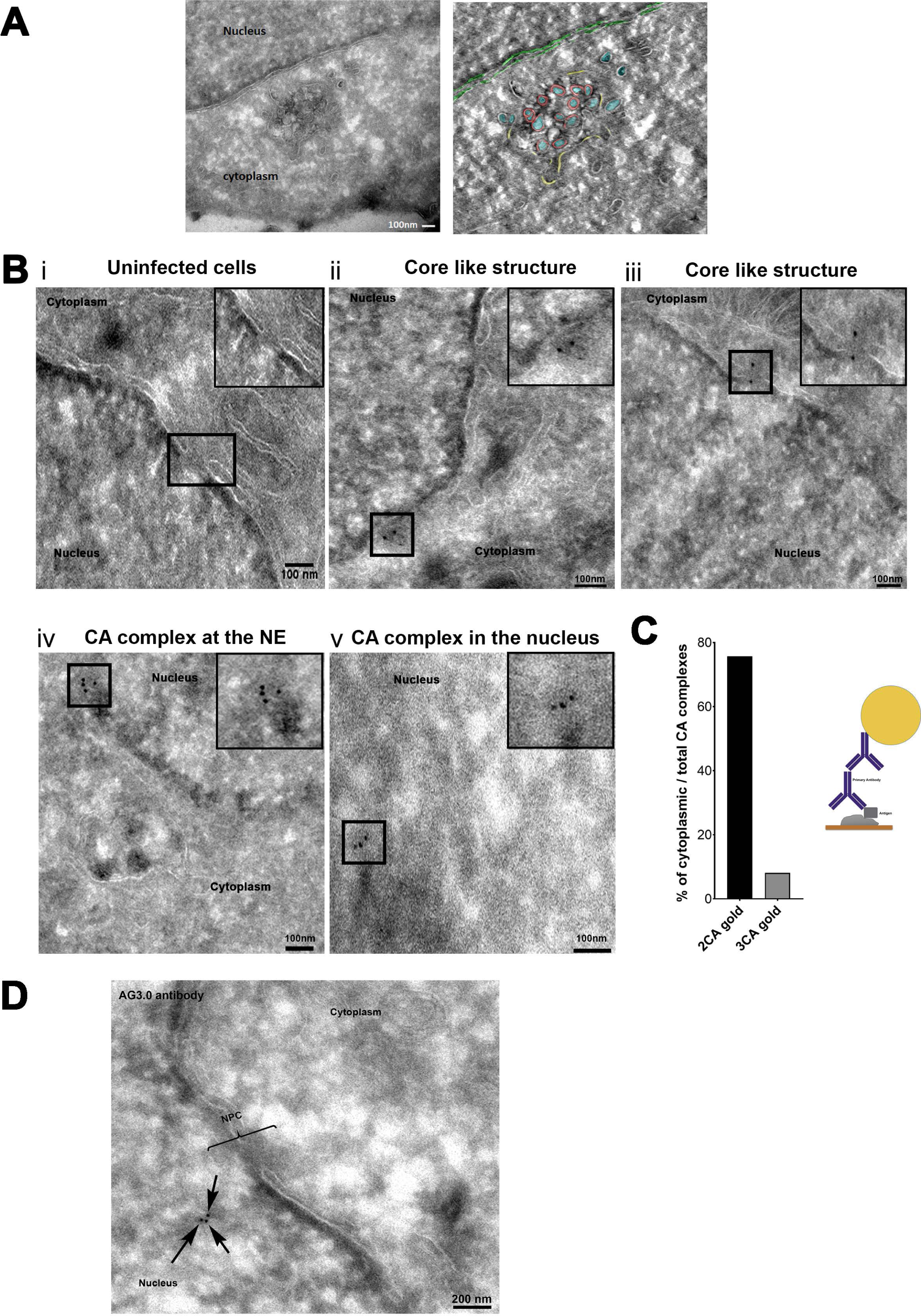
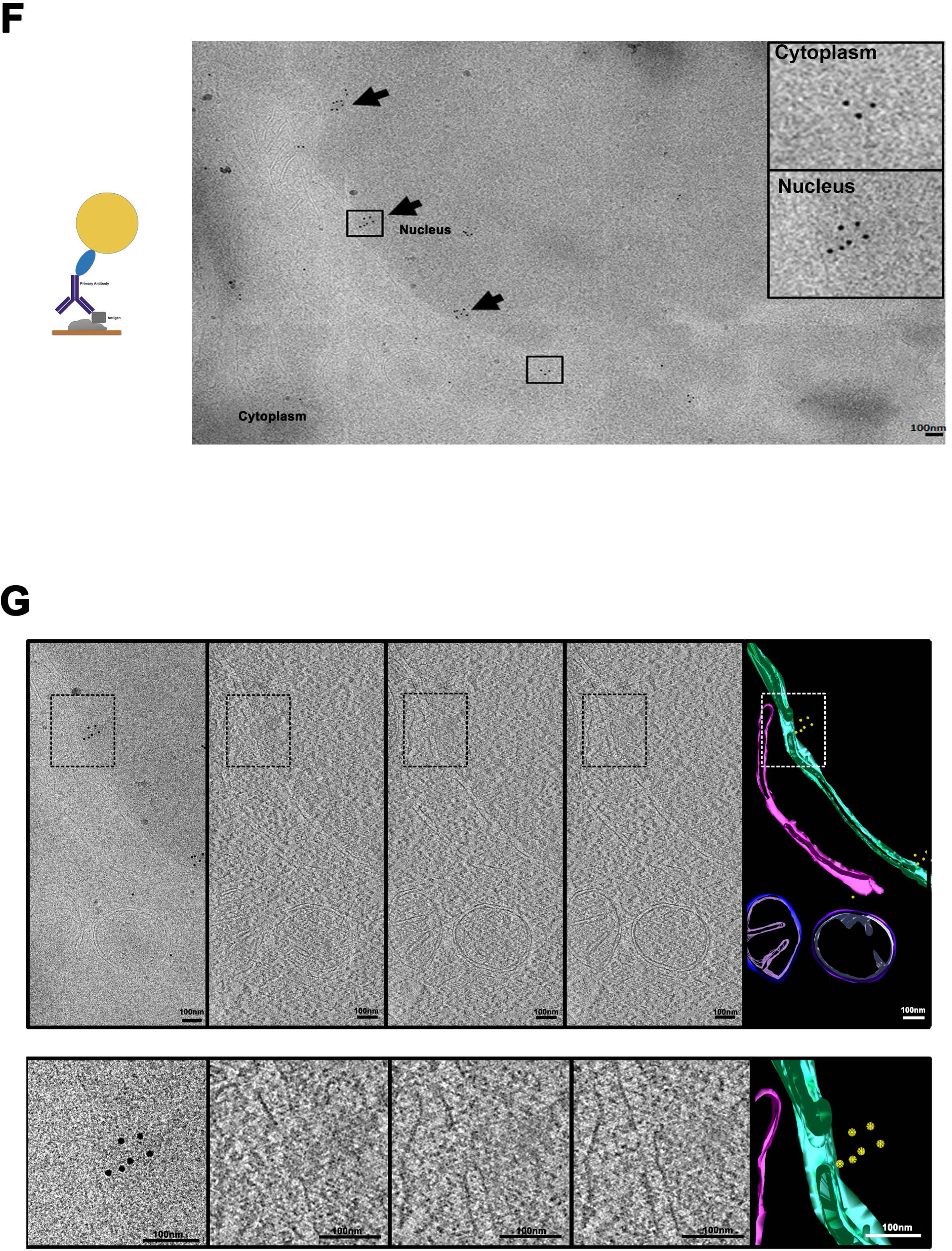
Remodelling of HIV-1 CA to translocate in the nucleus. (A) Images of HIV-1 virions englobed in the endosomes or cores escaping from the endosomes obtained by TEM. First from the left a 2D plane extracted from the tomogram of the endosomes containing viral particles, on the right the cell volume reconstructed and coloured (red for the envelope, blue for the cores, yellow for the borders of the endosome and green for the nuclear membrane). (B) Examples of structures resembling “core like structures” docked at the NPC by TEM. A negative control based on uninfected cells is shown in panel i. Panels ii and iii show “core like structures” docked at the NPC detected by 2 CA gold 10nm particles revealing the antibodies anti-p24 (NIH183-H12-5C) attached. Panels iv and v show intranuclear CA complexes. Scale Bar 100nm. (C) Quantification analysis of the CA gold particles distribution in viral complexes (∼ 39) located in the cytoplasm. The percentage of cytoplasmic CA complexes reveals that the majority of two gold particles (2CA) complexes are located in the cytoplasm contrary to the 3CA gold particles. Schema of the labelling method, first Ab anti CA and secondary Ab conjugated to gold particles. (D) HeLaP4R5 cells infected with HIV-1 IN_HA_ / VSVG. The sections were prepared and immunolabelled as in panel B, using a primary antibody anti CA (AG3.0). Black arrows indicate gold particles of 10 nm conjugated to a secondary Ab. Scale Bar 200nm. (E) HeLa P4R5 cells infected with HIV-1 IN_HA_ carrying on the WT envelope. The sections were prepared and immunolabelled as in panel B. The areas containing the viral CA complexes are enlarged in squares on the upper right of the pictures. Scale Bar 100nm. (F) Schema of the labelling method, first Ab anti CA and protein A gold. Cryo-electron and immunogold labelling have been used. The image contains several CA complexes (pointed out by black arrows) each one with multiple CA proteins showed by gold particles. The magnified views of the areas enclosed by black rectangles display the differences in the gold distribution between outside and inside the nucleus. Images where obtained using an antibody against HIV-1 CA followed by incubation with protein A coupled to 10nm gold. (G) Tomogram of a representative intranuclear CA complex highlighted with black rectangle in panel F. Sections were imaged in a T12 FEI electron microscope with tomography capabilities. The tomogram of the cell volume was reconstructed and manually segmented using IMOD. The upper panels starting from the left contain several planes (number 31, 43 and 49 out of 91 total slices) of the tomogram and the segmentation obtained from the reconstructed tomographic cell volume containing the gold labelled CA complexes, nuclear envelope (NE), endoplasmic reticulum (ER) and mitochondria (yellow, green, magenta and purple respectively). On the bottom is depicted the magnified views of the areas delimited by dashed lines on the upper panels.

**FIG. 3.**
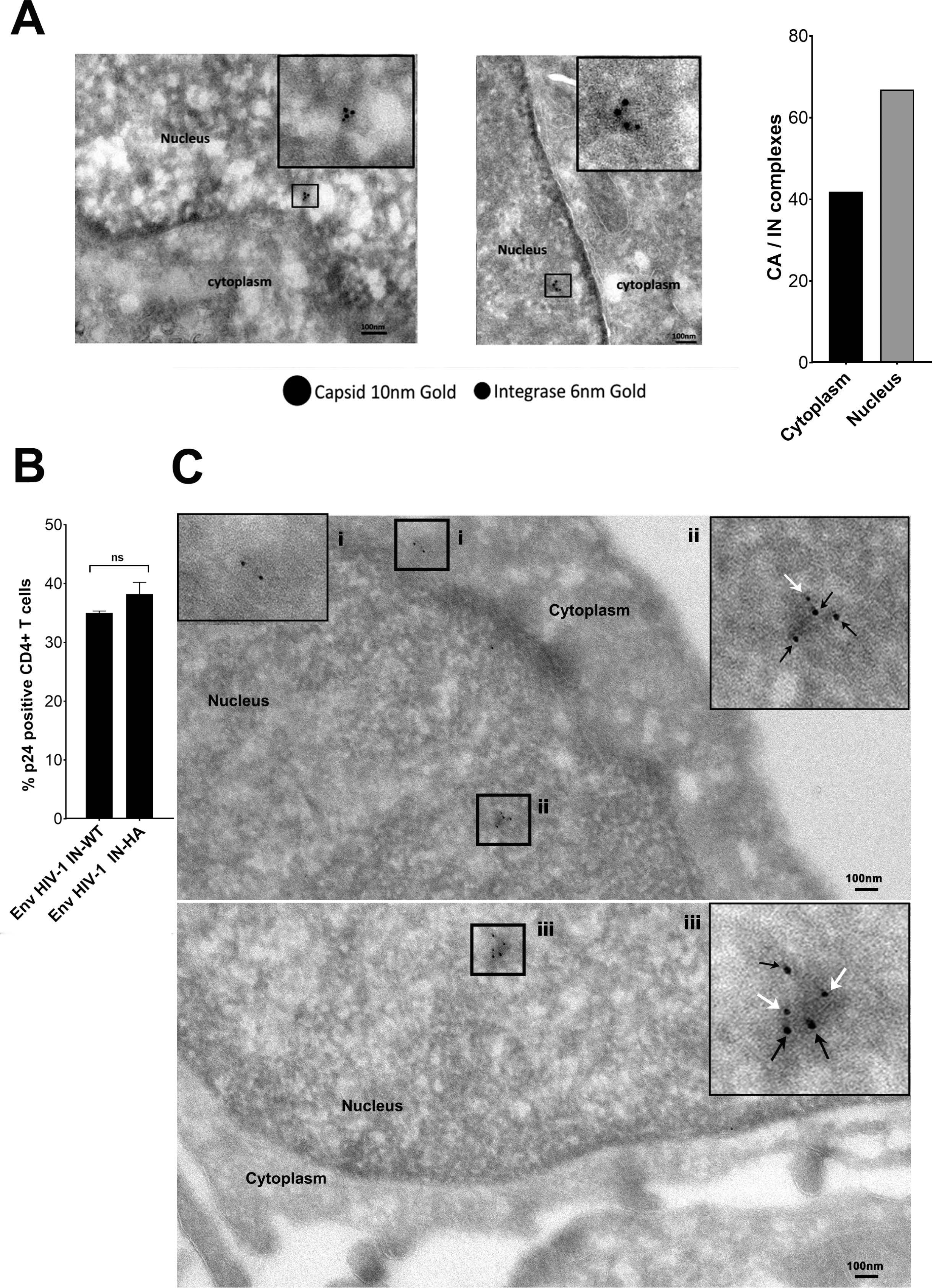
CA protein multimers are associated to IN inside the host nucleus. (A) Some CA complexes detected at 6 h post infection contain IN. The double labelling gold against primary antibodies anti-p24 (10nm) and anti-HA (to label IN) (6nm) highlights the association of both proteins particularly inside the nucleus of HeLa cells. CA/IN association in the cytoplasm and in the nucleus has been calculated and represented in the histogram on the right. B) Primary CD4^+^ T cells were isolated from healthy donors and infected with 4000ng of p24 of HIV-1. Comparison of the infectivity in primary CD4^+^ T cells (calculated as % of positive p24 cells analysed by cytofluorimetry) between wild type enveloped viruses carrying on the wild type IN or the IN fused to HA tag. C) CD4^+^ T cells were challenged with HIV-1 IN_HA_ / VSVG and samples were processed for immunoelectron microscopy after 9 hours from infection and stained with antibodies against capsid and HA for the integrase. Magnified insets represent: i) 2CA gold complexes located in the cytoplasmic area ii) and iii) 3CA complexes containing integrase as well. The black arrows correspond to capsid labelled with 10nm gold, the white arrows point the integrase labelled with 6nm gold.

Taken together, our data indicate that the viral core undergoes a morphological rearrangement during nuclear entry coinciding with an increased CA-labelling by EM likely due to increased accessibility of the antibody.

### Pearl necklace complexes contain IN and are present in the nucleus of CD4^+^ T cells

The presence of CA gold complexes in the nucleus of HeLa cells suggests that they are part of the viral complexes formed during nuclear translocation. Thus, to further characterize the composition and the spatial distribution of HIV-1 complexes inside and outside the nucleus, we labelled CA and IN with different sizes of colloidal gold conjugates (6 nm and 10 nm, respectively) and imaged them by TEM (Fig. 3A). IN was more frequently associated with intra-nuclear CA structures (Fig. 3A), that were detected by 3 or more CA gold rather than 2 CA gold particles. To investigate if these complexes could be observed also in natural HIV-1 target cells, we repeated the experiments in primary CD4^+^ T cells, the natural HIV-1 host cells (Fig. 3B). We observed viral complexes labelled by 2 gold particles in the cytoplasm of CD4^+^ T cells derived from healthy donors at 9 hours post-infection (Fig. 3C-i). The nuclear CA complexes detected by multiple gold particles were also observed in these cells (Fig. 3C-ii and -iii). Furthermore, the difference between the cytoplasmic and nuclear CA gold labelling was very similar in HeLa cells and in primary lymphocytes (Fig.2B and 3C).

Our results show that viral CA complexes associated with viral IN can be found in the nucleus of HeLa cells, and also in that of natural HIV-1 target cells, i.e. primary lymphocytes. Thus, the formation of the CA nuclear complexes was not cell dependent, implying that they represent an important feature for the nuclear journey of HIV-1.

### Specific detection of retrotranscribed DNA in infected cells by HIV-1 ANCHOR system

To investigate if the observed CA structures, detected during and after nuclear translocation, represent PICs, we analysed whether the retrotranscribed viral genome was present in these complexes. The rarity of viral events during nuclear import made their visualization particularly difficult. Labelling of the retrotranscribed viral DNA has been a major challenge and only partial success has been achieved using DNA FISH, View HIV assay, multiplex immunofluorescent cell-based detection of DNA, RNA and Protein (MICDDRP), or EdU in fixed cells (18, 29–31, 48–50). These DNA fluorescent labeling methods are all incompatible with TEM technique. This limitation impeded the characterization of the PIC during nuclear translocation. To overcome this drawback, we set up a new system that allowed direct tracking of viral retrotranscribed DNA in CA-positive complexes. We adapted ANCHOR technology (NeoVirTech) (51, 52) to visualize HIV-1 DNA (Fig. 4A). The ANCHOR technology consists in a bipartite system derived from a bacterial parABS chromosome segregation machinery. This is composed by a target DNA sequence, ANCH3, which is specifically recognized by the OR protein, a modified version of the bacterial parB protein (53, 54), fused to GFP. We cloned ANCH3 sequence in the HIV-1 genome (HIV-1 ANCH3) to ensure direct labelling of the retrotranscribed viral DNA in a highly specific manner (the human genome of the host cells lacks the ANCH3 sequences).We used this virus to infect HeLa cells previously transduced with a LV carrying OR-GFP cDNA (LV OR-GFP) (Fig. 4A and B). Our results by fluorescence microscopy revealed that HIV-1 ANCH3 is recognized by OR-GFP fusion proteins that accumulated on the target sequence resulting in the formation of a bright fluorescent spot (Fig. 4B). OR-GFP protein has no nuclear localization sequence (NLS) and therefore freely diffuses in the cell, with predominant localization in the cytoplasm (52). Upon infection, OR-GFP was efficiently visualized in complex with the retrotranscribed viral DNA, particularly in the nucleus (Fig. 4B). Of note, cells infected with HIV-1 ANCH3 or with the untagged HIV-1 showed similar number of proviruses by Alu PCR, thus both viruses behaved similarly during integration or steps of infection prior to integration (Fig. 4B). Importantly, HIV-1 ANCHOR permitted to track HIV-1 DNA by live-cell imaging. Hence, we could follow the fate of the viral DNA from the reverse transcription step onward. Indeed, this technology enabled us to follow the HIV-1 DNA for more than 70h in living cells by epifluorescence (Suppl. movies 1A,B), or by 3D or 2D spinning disk microscopy (suppl. movies 2A,B,C). Via the latter, we imaged infected cells to obtain complete information about the full cellular volumes. Next, to pinpoint the specificity of HIV-1 ANCHOR system to detect exclusively HIV-1 DNA, we infected HeLa OR-GFP cells with different MOIs (multiplicity of infection) of HIV-1 ANCH3. We observed a linear correlation between MOI and the number of nuclear vDNA spots in GFP+ infected cells (Pearson’s coefficient ∼ 1) (Fig.5A). The total number of intranuclear spots analysed for each condition was 2054 counts for 34 GFP+ infected cells (MOI 200,), 393 counts for 38 GFP+ cells (MOI 30), 290 counts for 44 GFP+ cells (MOI 10). Averages (Avg) of nuclear spots were calculated for single condition (MOI 10 Avg 6,7; MOI 30 Avg 10,07; MOI 200 Avg 60,4) (Fig. 5A). In addition, we infected cells in the presence of drugs, PF74 or nevirapine (NEV, inhibitor of RT). First we challenged HeLa cells expressing OR-GFP with HIV-1 ANCH3 for 24h without drug or in the presence of low and high doses of PF74 (Fig.5B and C). Both doses of PF74 blocked viral nuclear entry (Fig. 5D) (44). We detected the viral DNA inside the nucleus mainly in the absence of PF74 (Fig. 5B and C; suppl. movies 3A,B,C), in agreement with nuclear import data obtained by qPCR (Fig. 5D). Total intranuclear spots were analysed for each condition (no drugs 180 spots in 13 GFP+ cells; PF74 low dose 8 spots in 28 GFP+ cells; PF74 high dose 1spot in 27 GFP+ cells) (Fig. 5C). These results were confirmed also when the nevirapine was used. We counted intranuclear spots in 20 cells per condition and we obtained the following results: 152 nuclear spots in absence of NEV against 0 detections in presence of the drug. Thus, nuclear punctae containing HIV-1 DNA were found only in NEV untreated cells (Fig. 5E; suppl. movies 4 A,B). Overall, these observations demonstrated that HIV-1 ANCHOR technology faithfully tracked the retrotranscribed viral DNA (Fig. 5F).

**FIG. 4:**
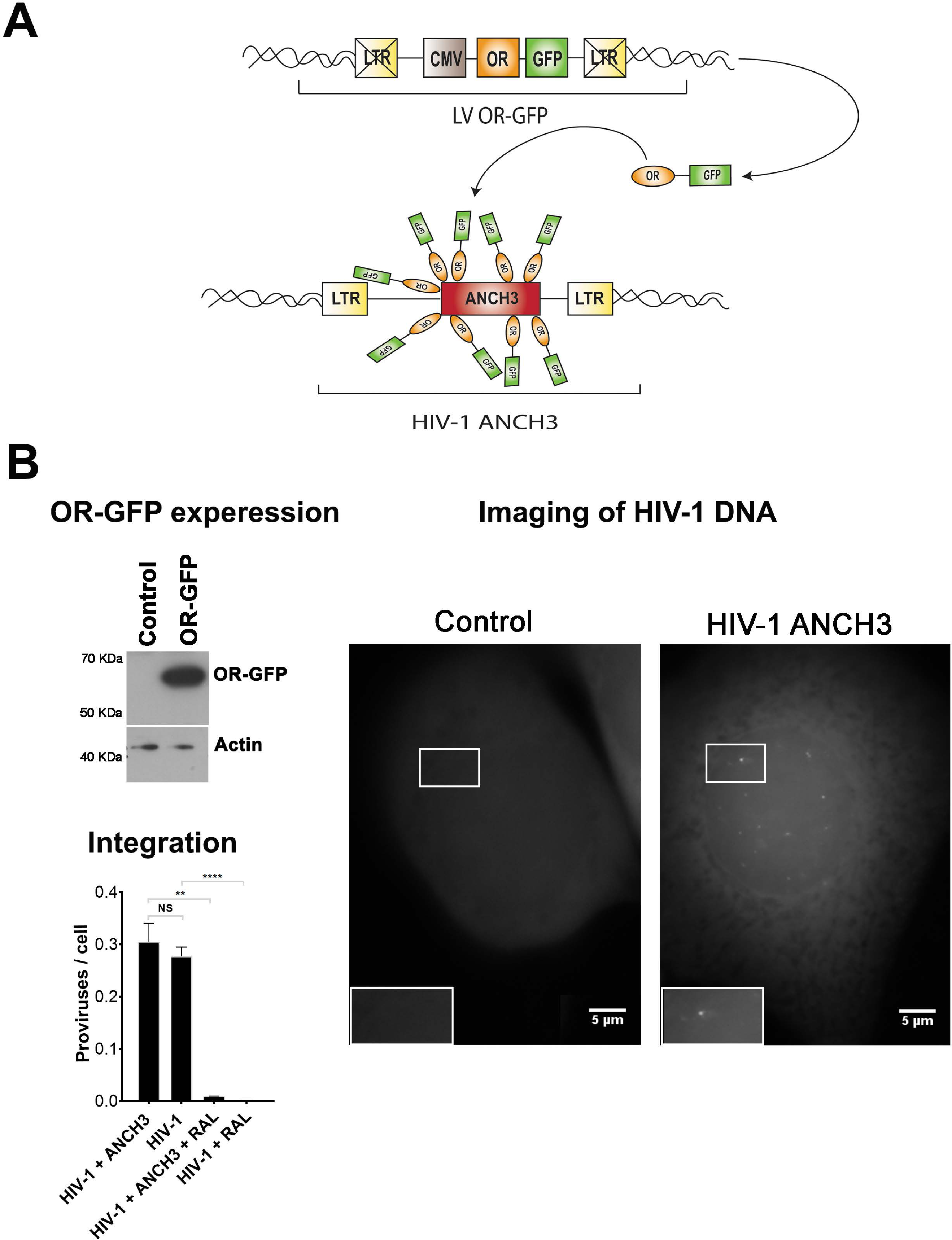
Detection of the retrotranscribed HIV-1 DNA in infected cells. (A) Schema of the HIV-1 ANCHOR system based on lentiviral vectors carrying on the OR-GFP cDNA under the control of CMV promoter (LV OR-GFP) and HIV-1 containing the ANCH3, target sequence of OR proteins (HIV-1ΔEnvIN_HA_ΔNefANCH3/VSV-G). (B) HeLa P4R5 cells were transduced with LV OR-GFP. The efficiency of OR-GFP expression was monitored by western blotting using antibody against GFP. As a loading control, samples were also blotted using antibody against actin. HeLa cells stably expressing OR-GFP infected or not with HIV-1 EnvIN_HA_ΔNef ANCH3/VSV-G (MOI 50) have been imaged by fluorescence microscopy at 24h post infection using water immersion objective in epifluorescence. The number of proviruses has been detected by ALU/PCR on HeLa P4R5 ORGFP infected with HIV-1 or HIV-1 ANCH3.

**FIG. 5.**
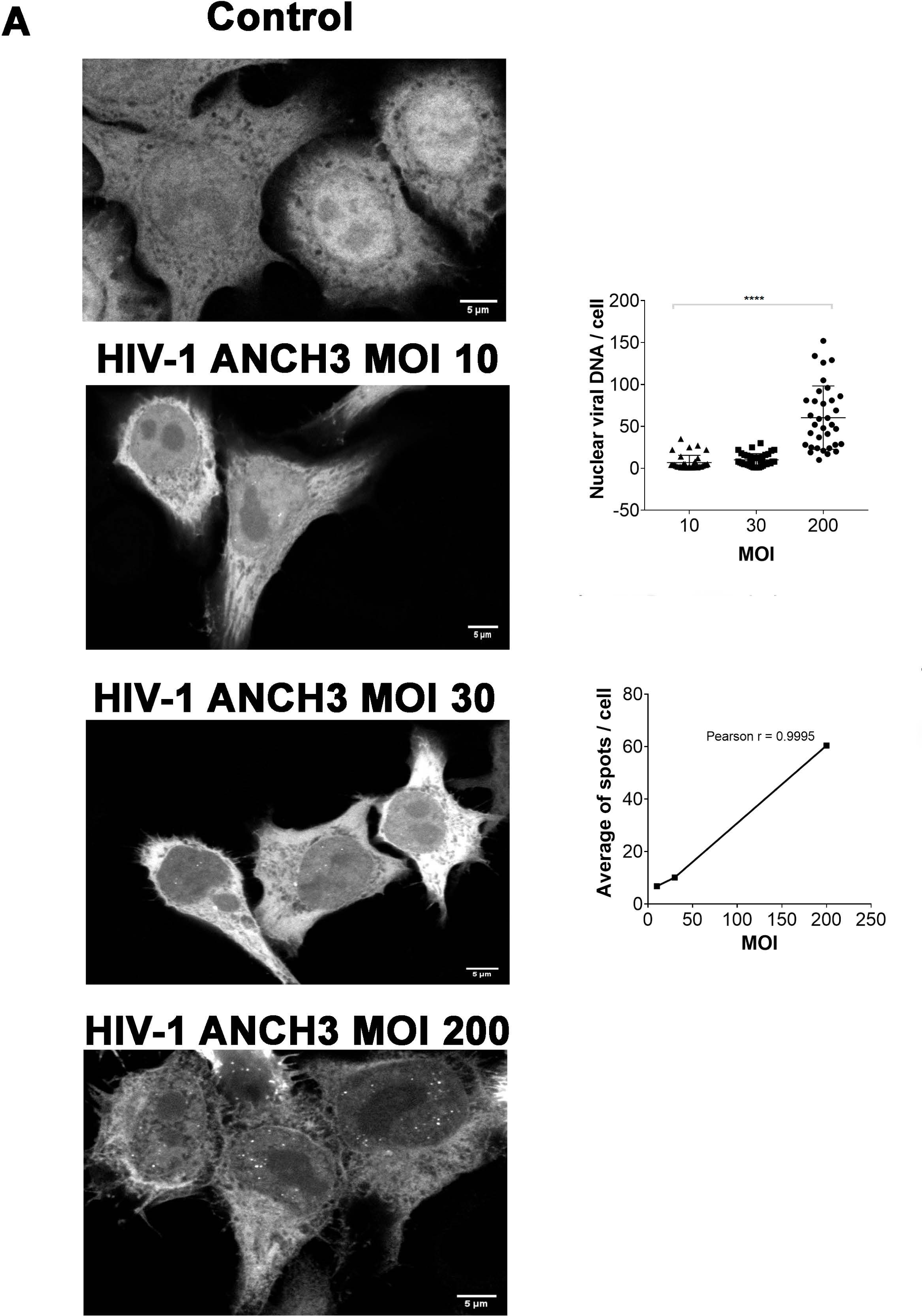
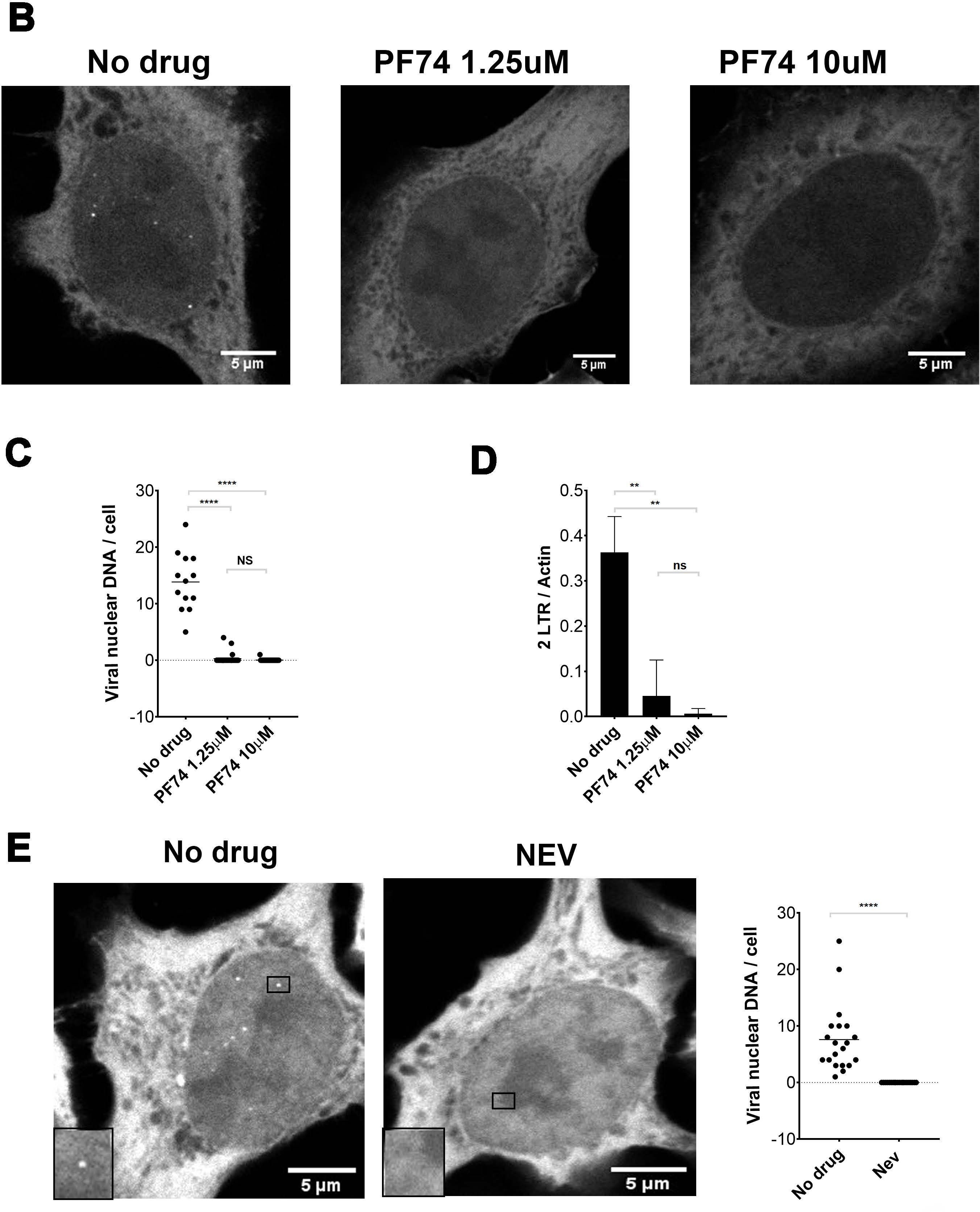
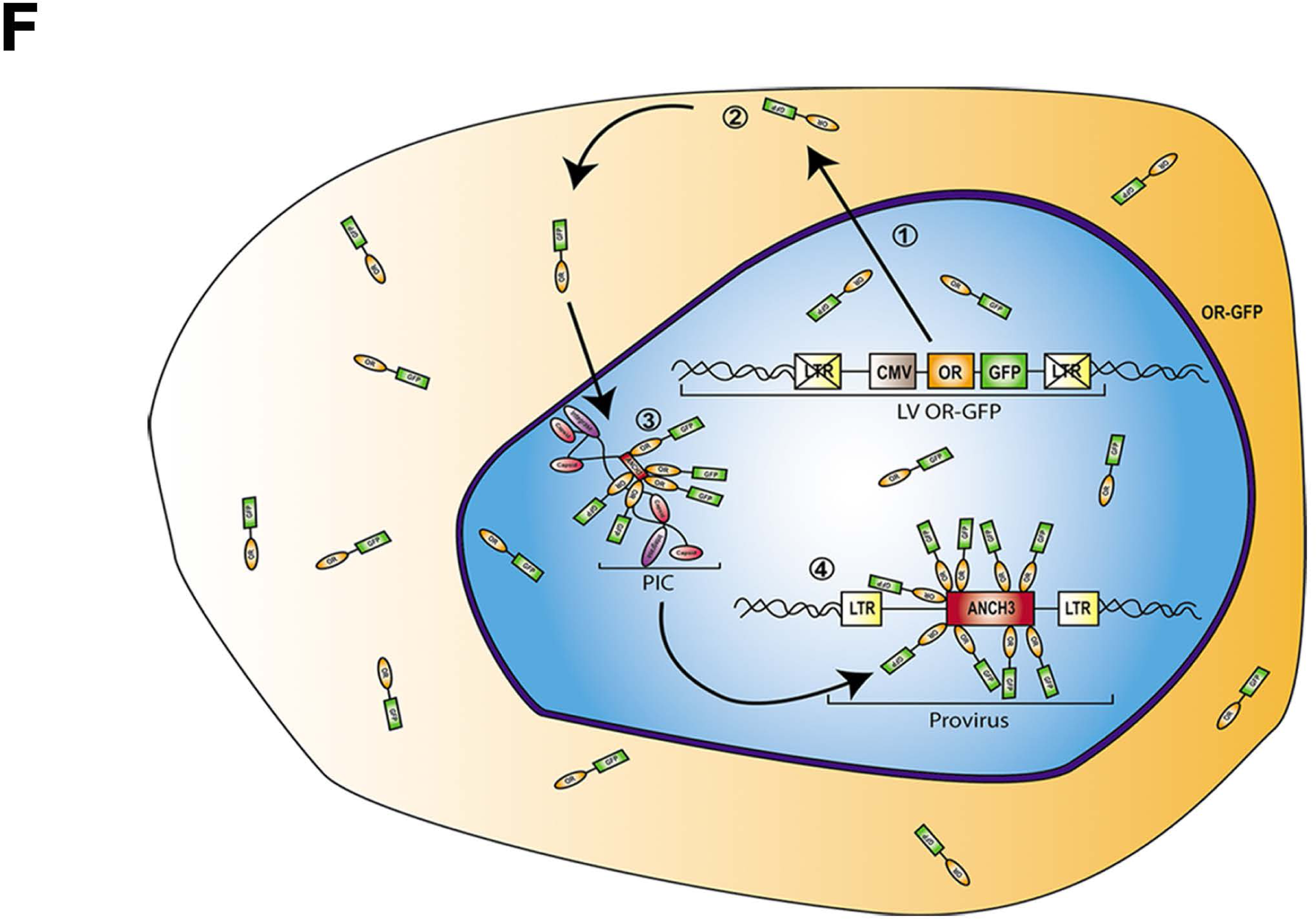
Specificity of HIV-1 ANCHOR system to detect the retrotranscribed DNA. (A) HeLaP4R5 cells stably transduced with LVOR-GFP were infected at different MOIs of HIV-1ANCH3 and imaged after 24 h by confocal microscopy. Nuclear viral DNA spots per single GFP+ cell were analysed in 2D by ImageJ. Correlation analysis and the Pearson’s coefficient as well as statistical analysis have been performed by Graph Pad Prism 7 (Anova test). (B) HeLaP4R5 cells infected at MOI 50 with HIV-1ΔEnvIN_HA_ΔNef ANCH3/VSV-G in presence or not of PF74 (low dose, 1.25µM; high dose 10 µM). Cells were imaged by confocal microscope at 24h post-infection. (C) Individual spots inside the nuclei were manually counted and statistically analysed in 2D by Graph Pad Prism 7, statistics were calculated using two-tailed Student’s t test, P value <0.0001 (****) and nonsignificant (ns). (D) Viral nuclear import has been evaluated by qPCR (2LTRs) and normalized by actin. Statistical analysis has been calculated by Graph Pad Prism 7 using two-tailed Student’s t test. Differences were considered statistically significant at a P value of <0.001 (***), or <0.01 (**).(E) Confocal microscopy of intranuclear spots detections in HeLa P4R5 ORGFP challenged with MOI30 of HIV-1 ANCH3 in presence or not of NEV at 24h post-infection. 2D statistical analysis of a manual count of intranuclear spots has been performed by Graph Pad Prism 7. All data are representative of two or more independent experiments. (F) Schema of the HIV-1 ANCHOR technology used to live-track the viral DNA. 1) The expression of OR-GFP fusion protein by LV OR-GFP; 2) OR-GFP is translated in the cytoplasm and diffuses in the whole cell volume with a main location in the cytoplasm due to the lack of the NLS; 3) binding of OR-GFP to ANCH3 sequence contained in the incoming PICs 4) or in the nuclear viral DNA forms.

### HIV-1 CA decorates the retrotranscribed viral DNA during nuclear translocation

Our ultimate goal was to characterize the CA- and vDNA-positive complexes during and after nuclear translocation. One of the advantages of the OR-GFP system is its compatibility with TEM. Since this system is based on the interaction of OR protein with the ANCH3 sequence we were able to apply a correlative light and electron microscopy approach. Thus, we coupled HIV-1 ANCHOR to immunogold labelling to investigate if the viral DNA was part of the “pearl necklace” CA shapes previously detected by TEM (Fig. 2B-iv,-v; 2D, F and G; 3A and C-ii,-iii). We transduced HeLa cells with LV OR-GFP and 3 days later we challenged those cells with HIV-1 ANCH3 for 6 hours. Cells were fixed and prepared for EM-immuno-labelling to test if structures positive for CA also contained viral DNA. Hence, we observed HIV-1 CA complexes near the nuclear pore complex (NPC) at 6h post infection. These CA complexes were positive for DNA as observed by the correlative method (Fig. 6A). CA-positive complexes typically decorated by 3 CA gold particles, were also associated with HIV-1 DNA during nuclear translocation (Fig. 6B). The error of correlation between the two images (EM and fluorescence) has been calculated by ecCLEM plugin on Icy software (55, 56) to be theoretically ∼ 70nm. CLEM results show a more elongated CA-positive form during nuclear translocation (Fig. 6B) than the “core like shape” observed in the cytoplasm (Fig. 2A and B-ii,-iii). These results strongly suggest in which form a viral complex can fit the size of the NPC and translocate through the pore to reach the host nuclear environment. Additionally, we performed a dual gold labelling experiment to detect viral complexes inside the nucleus. We used different size of gold particles to label the viral DNA through OR-GFP (anti-GFP, 5 nm gold) and the viral CA (10 nm gold). Thus, we were able to reveal complexes formed by the viral DNA associated to HIV-1 CA in the nucleus of infected dividing cells (Fig. 7). These data are in line with our CLEM results, showing that viral complexes containing the retrotranscribed DNA can retain several CA proteins even after nuclear translocation.

**FIG. 6.**
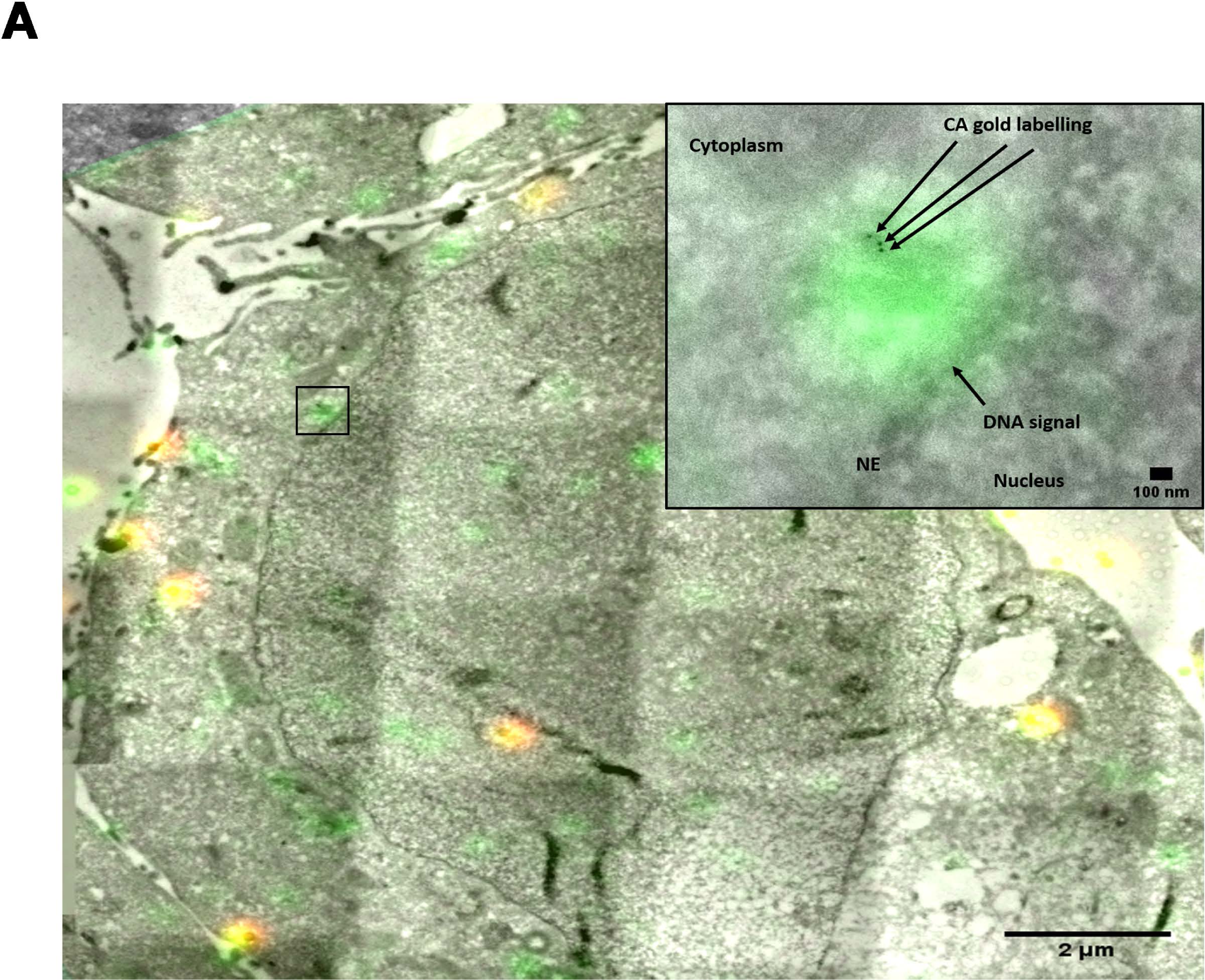
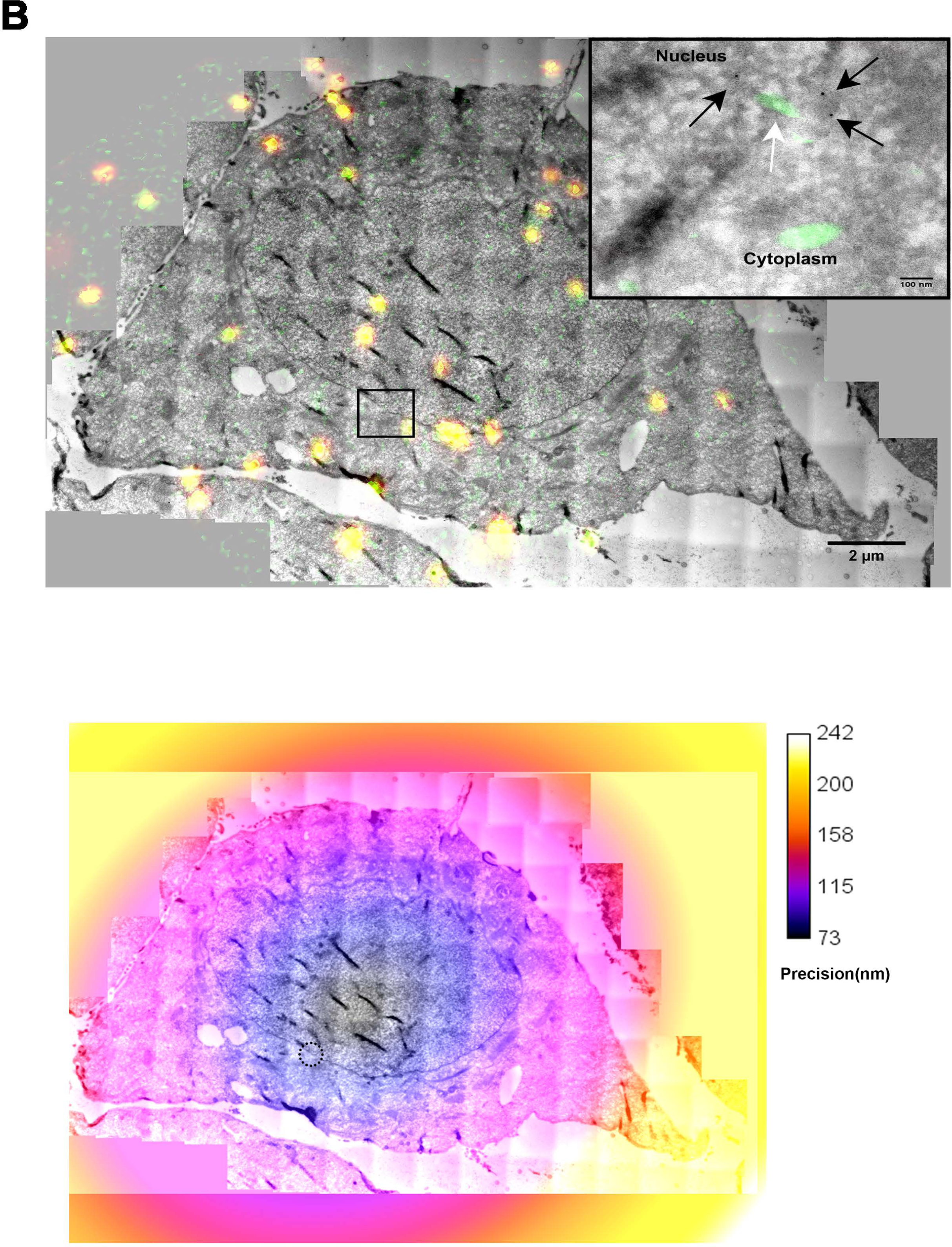
Multiple CA proteins leading retrotranscribed DNA detected by CLEM. (A) HeLa P4R5 transduced with OR-GFP were infected with HIV-1 ANCH3 and processed for correlative light and electron microscopy (CLEM). To achieve a precision correlation, fluorescent beads of 200nm were added, visible also by EM. CLEM results on HeLa P4R5 OR-GFP cells at 6 h p.i. show the GFP signal in green revealing the location of HIV-1 DNA. The signal of OR-GFP is amplified by using an antibody against GFP. HIV-1 CA is detected by a first antibody anti-CA and a secondary antibody gold conjugated (gold 10nm). The yellow signals correspond to the beads emitting in green and in red channels. On the top right, the magnified view of the black square is shown. The green signal shows the location of viral DNA, associated to the CA gold labels, dark dots. (B) Another biological replicate of CLEM shows multimer of CA proteins (dark dots) shrouding the vDNA (green signal) during nuclear translocation. Panel on the bottom shows an example of the precision correlation process applied. The precision of the correlation between TEM and fluorescence images were estimated with ec-CLEM plugin under the icy environment. The calibration bar represents the precision achieved in nm by the different area of the cells. The dashed circle shows the area enlarged in the black box of the top panel.

**FIG. 7.**
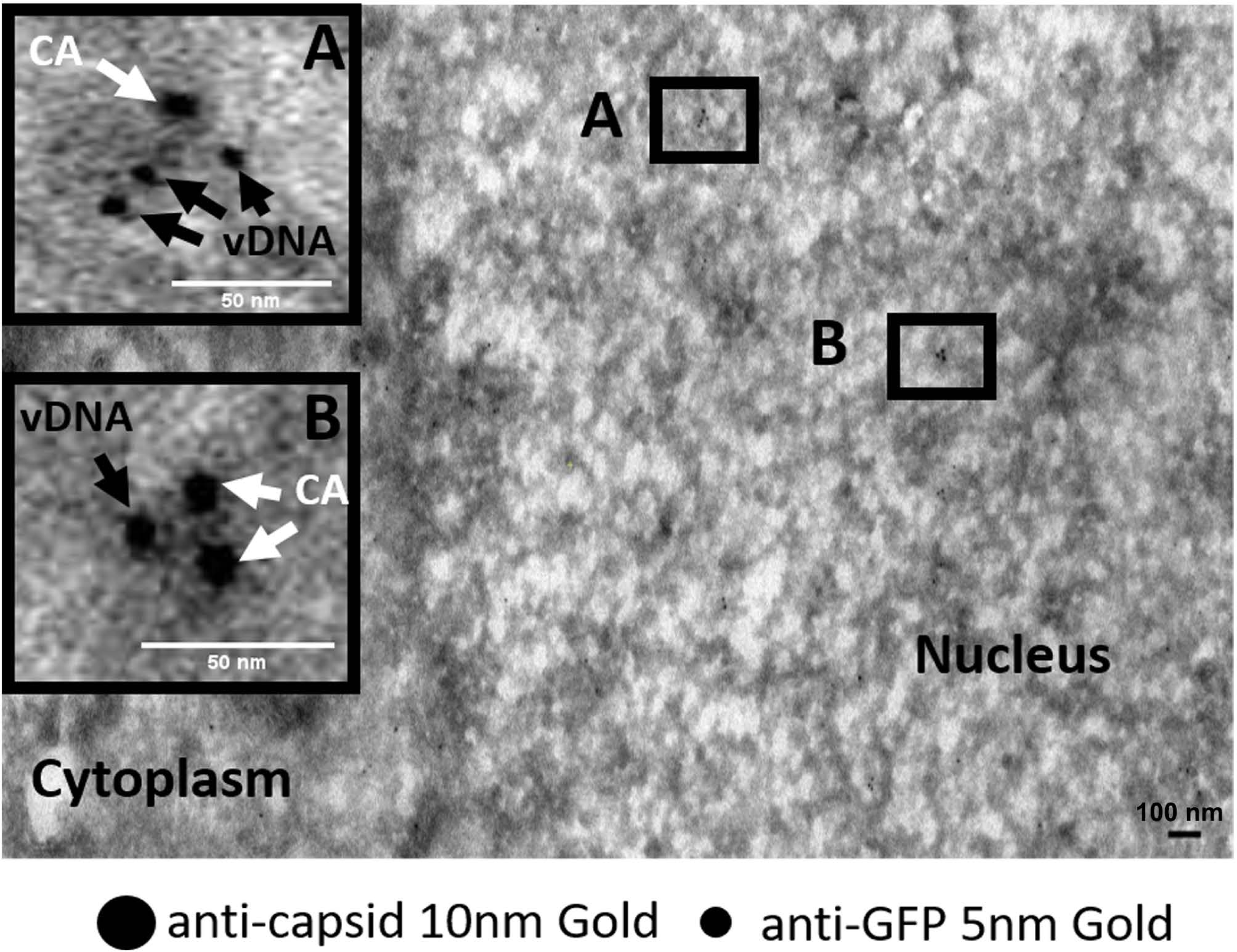
Intranuclear viral complexes. Double gold labelling coupled to TEM show CA / OR-GFP (viral DNA) as part of the same intranuclear complexes. Viral DNA is detected by the presence of clusters formed by multiple OR-GFP bound to ANCH sequence cloned in HIV-1 genome. OR-GFP proteins are labelled by the same primary antibody against GFP used in CLEM and a secondary antibody conjugated with gold particles of 5nm. HIV-1 CA is revealed by a primary antibody against CA (NIH183-H12-5C) and a secondary antibody conjugated with gold (10nm). Scale bars 100nm.

Overall results obtained by TEM and by CLEM highlighted the shape of HIV-1 PIC during and after nuclear entry, including all required components for the integration, such as the integrase, DNA and, surprisingly, capsid forms composed by multiple CA proteins.

## Discussion

HIV-1 core must undergo structural changes to be able to cross the NPC, as the pore is too narrow to allow its passage as an intact core. Thus, the organization of viral CA complexes during the early steps of HIV-1 infection is still unknown, due to the lack of appropriate tools to study this event. Our study provides some new evidences on the viral remodelling of HIV-1 core occurring prior, during and after viral nuclear entry. Importantly we confirmed such observation in the main physiological target cell of HIV-1, CD4+ T cells. We were able to quantify a different distribution of CA gold labelling comparing cytoplasmic and nuclear structures (Fig. 2C), supporting a viral CA remodelling process during nuclear entry. We observed that multilobe structures resembling to “core like shapes” can be labelled by 2 gold particles, although the HIV-1 core is composed by ∼1500 CA monomers (12, 57). This can be explained by specific features of EM immuno-labelling that is dependent on the specificity of the antibody, accessibility and number of antigens on thin sections, amongst other factors (58). Intranuclear CA complexes were detected by more gold particles than the cytoplasmic viral structures, which we believe is due to rearrangement of the viral CA (Fig. 2B-iv,-v, F and G). It should be noted that so far the presence of multiple CA proteins inside the nucleus of infected mitotic cells has never been reported. To confirm the detection of intranuclear complexes, tomograms performed at 6 hours post infection revealed the formation of pearl necklace shapes composed by multiple CA located in the nucleus (Fig. 2B-iv,-v, F and G). Often these nuclear CA structures are associated to viral IN (Fig. 3A). Importantly, similar complexes composed by multiple CA and IN have been identified in HeLa cells and primary lymphocytes, revealing the organization of the viral CA proteins also in the main target cells of HIV-1 (Fig. 3C). Our results show that viral complexes observed during nuclear translocation could represent PICs, thanks to the ability to combine the specific labelling of the retrotranscribed DNA (HIV-1 ANCHOR) with immunogold CA labelling. So far the viral DNA has never been visualized associated to multiple CA proteins in dividing infected cells. Importantly, we observed that HIV-1 ANCHOR allowed the specific detection of the viral DNA as shown by the use of drugs (Fig. 5). This is highlighted by using drugs that block different steps of the HIV-1 early events, such as nuclear import (PF74 drug) or reverse transcription (nevirapine) (Fig 5). Bright spots identifying viral DNA were prevalently detected in the nucleus of untreated cells, in agreement with results obtained by real time PCR that amplified 2LTR circles (Fig. 5B, C and D), commonly used to detect viral nuclear import (59). Thus, CLEM enabled to visualize the organization of HIV-1 PIC during nuclear translocation in HIV-1-permissive cells. It should be noted that HIV-1 ANCHOR can specifically label all nuclear viral DNA forms, thus, further experiments are needed to distinguish integration competent viral complexes from dead-end products. Independently of the fate of the PIC, HIV-1 ANCHOR technology combined with EM offered the opportunity to observe how viral CA proteins reshape near the NPC to lead the viral genome in the nucleus (Fig. 6A and B). OR-GFP can bind the viral DNA only if it is exposed, as indicated by our CLEM results, which support our model of viral core remodelling. In fact, complexes of multiple CA associated to the retrotranscribed DNA can be visualized by CLEM before and during the nuclear translocation (Fig. 6A and B). Likely, the observed complexes contain complete retrotranscribed genomes. In agreement with this observation, previous studies showed that OR-GFP binds only dsDNA (60), meaning that the vDNA shrouded in the multiple CA proteins complex is a dsDNA. The ANCH3 sequence has been cloned in place of nef, which means that the vDNA identified by CLEM might be a late reverse transcript, since OR-GFP can only bind it after the completion of reverse transcription on both strands containing the ANCH3 sequence. Thus, all viral complexes carrying an accessible vDNA can be observed by fluorescence microscopy. As opposed to fluorescence microscopy, the visualization of viral complexes by EM coupled with gold labeling is more complicated, but has the advantage of yielding high resolution images. Sections are not permeabilized, so only CA epitopes exposed on the surface of the section can be labeled by the anti-CA antibody and then by a secondary gold labeled antibody. Indeed, a limited number of available epitopes can be recognized by the primary antibody in sections. The aforementioned reasons, together with the thorough sample fixation required for the EM strictly limit the number of epitopes accessible to the antibody. Apart from such technical limitations, we do not expect to have all vDNA complexes co-labeled with CA, as a result of the asynchronous infection conditions used and the consequent presence of some CA-free nuclear vDNA, as previously shown by Brass group (30). Overall, these results indicate that the viral complexes visualized by CLEM could potentially be PICs.

Summarizing, the viral CA reshapes before crossing the NPC, decorating the viral DNA and leading it into the nucleus. Similar CA-viral DNA complexes can be also found inside the host nuclei (Fig. 7).

The presence of consistent shapes formed by multiple CA proteins in the nucleus (Fig. 2B-iv,-v, D, F and G; Fig. 3A and C), as well as their association to the retrotranscribed DNA (Fig. 6A and B; Fig. 7), would indicate that these CA forms are imported with the viral genome into the host nucleus. These results would support the evolution of the concept of viral uncoating, no more seen as a complete loss of viral CA but as a CA remodelling during nuclear import. Another novel aspect of our work is the development of the HIV-1 ANCHOR system, the first fluorescence DNA labelling technique shown to be compatible with TEM. This innovative technology allows us to follow the HIV-1 DNA shrouded by multiple CA proteins during and after nuclear entry in mitotic cells. In addition, we were able to live-track the viral DNA in the nucleus of infected cells following translocation through the pore (suppl. movies 1,2,3,4).

Of note, our study gives new information on the early steps of HIV-1 infection in dividing cells. It is known that mitotic cells with integrated viruses may persist for many years, undergo clonal expansion (61). Clonal expansion seems to be the major mechanism to generate HIV-1 reservoir (62), considered to be established early during primary HIV-1 infection (63).

Overall, our results provide new notions not only on the organization of viral PIC complexes but also on the dynamics and the fate of the viral DNA inside the nucleus. Our data elucidate how the viral CA remodels to lead HIV-1 DNA in the nucleus of infected cells.

These findings and technology can be useful for future studies on other pathogens or to investigate the interplay of HIV-1 DNA with nuclear factors and chromatin environments.

## Methods

### Cells

HeLaP4R5 cells, a HeLa-CD4/LTR-lacZ indicator cell line expressing both CXCR4 and CCR5, were employed to assess viral infectivity (64) using a beta gal assay. 293T cells (ATCC) are human embryonic kidney cells used to produce lentiviral vectors and HIV-1 viruses, HeLa cells (ATCC) derived from cervical cancer cells. CD4^+^ T cells were isolated from healthy donors obtained via the EFS (Etablissement Français du Sang, Paris, France). Briefly, primary CD4^+^ T cells were purified from human peripheral blood by density gradient centrifugation (Lymphocytes separation medium, PAA) followed by positive immunomagnetic selection with CD4 Microbeads (Miltenyi). The day later cells were activated by T Cell Activation kit (Miltenyi) for 2-3 days at 37°C with interleukin 2 (IL-2)-containing medium (50 IU/ml), then cells were challenged with HIV-1 carrying the CA wild type or mutated. Percentage of p24 positive cells has been obtained cytofluorimetry acquisition and FlowJo analysis.

### Antibodies

Ab anti-actin HRP conjugated sc-2357 Santa Cruz (dil. 1:5000), Ab anti-p24 antibody NIH183-H12-5C or AG3.0 (NIH reagent, IF dil. 1:400 or TEM 1:50) and the anti-HA high affinity antibody (11867423001) Roche (TEM 1:50 dilution or IF 1:500), Ab Goat anti-mouse Alexa Fluor Plus 488 (A32723) and Goat anti-rat Alexa 647 (A21247) Thermofisher scientific. Ab Goat anti-mouse 10nm gold coupled (ab39619), Ab Goat anti-rat 6nm gold coupled (ab105300) Abcam (dil. 1:50), Ab Goat anti-rabbit coupled to 5nm gold Abcam (ab27235). Ab anti-GFP rabbit (ab183734) Abcam (CLEM dil. 1:50), Ab anti-GFP (Clontech #632592, WB dilution 1:1000), Ab Beta Actin HRP conjugated (Abcam, #8226 WB dil. 1:2,500), Ab Goat anti-rabbit Alexa 488 (A11078) (CLEM dil.1:50), Ab anti Nup153 9 (kind gift from B. Burke dil. 1:200), Protein A-10 nm gold (dil. 1:50) from UMC-Utrecht, Netherlands

### Time-lapse microscopy

HeLaP4R5 cells stably transduced with LV OR-GFP were plated in HiCQ4 microdishes (10,000 cells per chamber) (Ibidi). The following day, cells were infected with HIV-1ΔEnvIN_HA_ΔNefANCH3/VSV-G. Transmission and fluorescence images were taken every 5 or 10 min for up to 96 h using a Nikon Biostation IMQ (40X objective) with 6-8 fields captured simultaneously for each condition or for up to 24h by using a spinning-disk UltraView VOX (Perkin-Elmer) (63x objective) with one field of view for each experiment in 2D or 3D. Images were analyzed in FIJI or Imaris.

### Western blotting and confocal immunoimmunofluorescence microscopy

The expression of the correct size of the cDNA OR-GFP cloned in LV has been tested by western blotting. Proteins were extracted on ice from wild type and LVOR-GFP transduced HeLa cells using RIPA buffer (20mM HEPES pH 7.6, 150mM NaCl, 1% sodium deoxycholate, 1% Nonidet P-40, 0.1% SDS, 2mM EDTA, complete protease inhibitor (Roche Diagnostics)), and protein concentration was quantified using the Dc Protein Assay (Bio-Rad Laboratories) with BSA as standard. Ten micrograms of total protein lysate was loaded onto SDS–PAGE 4–12% Bis Tris gels (Invitrogen). Revelation was carried out using the ECL Plus western blotting kit (GE Healthcare). Primary antibody used for western blotting (WB) was anti-GFP (Clontech #632592, dilution 1:1000). Secondary conjugated antibodies used for western blotting were Beta Actin HRP conjugated antibody (Abcam, #8226 1:2,500), and anti-rabbit IgG HRP (sc2357 Santa Cruz). Immunofluorescence microscopy: HeLa P4R5 cells stably expressing OR-GFP or not were plated onto 12 mm diameter coverslips in 24-well plates the day before and then infected with HIV-1ΔEnvIN_HA_ΔNef ANCH3/VSV-G or HIV-1ΔEnvIN_HA_ /VSV-G at different MOIs and different time post infection. The cells were then washed, fixed with 4% PFA, permeabilized with Triton X 100 0.5% for 30 min and blocked with 0.3% bovine serum albumin (BSA). All incubations were carried out at room temperature and were followed by five PBS washes. Cells were incubated with primary antibodies for 1 h and secondary antibodies for 30 min. Antibodies were diluted in 0.3% BSA. Nuclei were stained with Hoechst (Invitrogen, dilution 1:10000). Finally, cells were mounted onto glass slides (Thermo Scientific) with Prolong Diamond (Life Technologies). Confocal microscopy was carried out on a Zeiss LSM700 using a 63× objective. Representative medial sections or combined Z stacks are shown as indicated. Images were analyzed in FIJI.

### Viral infection and sample preparation for electron microscopy

Eight million of HeLa P4R5 or Hela P4R5 OR-GFP transduced cells were seeded in a T75 flask and infected with 4000ng of p24 of either HIV-1 IN-HA or HIV-1 ANCH3 and incubated for 6h. When a WT virus has been used to infect HeLa P4R5 cells or primary CD4^+^ T cells a ultracentrifuged virus has been used in some cases in presence of SEVI according a published protocol (65, 66) (SEVI fibrils have been kindly provided by Franck Kirchhoff). Infectivity has been analysed by beta gal assay or by FACS. Samples were prepared for EM as follows: cells were fixed by adding directly an equal volume of 8% paraformaldehyde, 0.2% glutaraldehyde in PHEM buffer (60mM Pipes, 25mM Hepes, 2mM MgCl_2_, 10mM EGTA, pH 7.3) solution to the cells medium and incubated for 30 minutes. Next, the solution was exchanged by 4% paraformaldehyde diluted in PHEM buffer and incubated for 2 hours at room temperature. Cells were further prepared for cryomicrotomy and immunolabelled as described in(67). Electron microscopy chemicals were purchased from Electron Microscopy Sciences (Pennsylvania). For the CLEM experiments before contrasting with uranyl acetate the samples were stained with Hoechst 1µM for 20 minutes in water, washed and incubated with a solution of 0.2µm Tetraspeck fluorescent beads (Thermofisher scientific) diluted 1:50 in PHEM buffer pH 7.3 for 20 minutes and washed 4 times 2 minutes with water. The samples were mounted on in a glass bottom petri dish (Miltenyi Biotec) with a drop of SlowFade Diamond antifade mountant (Thermofisher Scientific). The imaging process gave a mosaic map of the sections in the blue, green and far red channels using a 63X 1.4 NA objective with a Leica DSM6000 microscope equipped with Orca Flash 4.0 LT camera (Hamamatsu Photonics). Then the grids were recovered by pouring 10ul of water underneath them. Grids were washed contrasted and prepared for TEM as specified above. For the cryo-EM observation the samples were prepared as described above. After immunolabelling the grids were embedded with a mixture of 50% methylcellulose 2% and 50% sucrose 2.3M and then vitrified by plunge freezing with EMGP plunge freezer (Leica) at 30 °C and 90% humidity.

### Electron microscopy data collection and image processing

Sections, at room temperature or in cryo, were transferred and imaged in a Tecnai T12 transmission EM operating at 120 or 80kV equipped with a Gatan Ultrascan 4000 camera. Multiscale mapping and tilt series acquisitions in areas of interest were processed by a Serial EM software(68). In case of cryo samples, low dose conditions and bi-directional tilt schemes were used during acquisition. Tilt series stacks were initially aligned using cross-correlation and the alignments were further refined using the immunogold beads as registration fiducials in IMOD (69). Tomograms were reconstructed with the weighted back-projection method and filtered to assist manual segmentation with IMOD.

The correlation between fluorescence and electron microscopy were achieved using the following protocol: 1) z-stacks of every frame of the mosaic was projected with the FIJI’s plugin extended depth of field(70); 2) the frames were aligned and blended to generate a fluorescence map of the complete section using Mosaic J (71); 3) the same cells was identified in both fluorescence and low resolution TEM section map; 4) the high precision correlation was obtained by identifying Tetraspecks positions in high resolution fluorescence and TEM images using ec-CLEM plugin (56) of Icy (55).).

### Quantitative PCR

Total cellular DNA was isolated using the QIAamp DNA micro kit (QIAGEN) at 7 and 24 h post infection. The genomic DNA was treated for 1h at 37 °C with Dpn1. Ten micromolar of nevirapine was used in infected cells as control of the experiment. Late reverse transcription products at 7 h post infection were measured by real-time PCR using primers and probe previously described (59), 2LTR containing circles were detected using primers MH535/536 and probe MH603, using as standard curve the pUC2LTR plasmid, which contains the HIV-1 2LTR junction. Integration was assessed by Alu-PCR, using primers designed in the U3 region of LTR (4) which is deleted in the LVs carrying OR-GFP but not in the LTR of HIV-1 used to challenge OR-GFP stably expressing cells and control cells. The standard curve has been prepared as follows: DNA generated from infected cells was end point diluted in DNA prepared from uninfected cells and serial dilutions were made starting from 50,000 infected cells. Each sample amplified contained 10,000 infected cells mixed at 40,000 uninfected cells. The control of the first round PCR was the amplification without Alu primers but only U3 primers (4). Dilutions 1:10 of the first round were amplified by real time PCR (4). Internal controls have been included such as infection in presence of RAL (10µM). LRT, 2-LTR and Alu-PCR reactions were normalized by amplification of the housekeeping gene Actin (4).

### Plasmids and viral production

HIV-1ΔEnvIN_HA_ΔNef ANCH3 plasmid was obtained by insertional mutagenesis using Quik Change II XL Site-Directed Mutagenesis kit and the sequence ANCH3 has been cloned by PCR using as template the plasmid pANCH3 (NeoVirTech). The ANCHOR^TM^ technology is the exclusive property of NVT. The LVCMVOR-GFP was generated by cloning by PCR OR-GFP from the plasmid pOR-GFP (NeoVirTech) in pTripCMVGFP. Lentiviral vectors and HIV-1 viruses were produced by transient transfection of 293T cells using calcium phosphate coprecipitation. Lentiviral vectors were produced by co-transfection of 10 µg of transfer vector LV OR-GFP with 2.5 µg of pMD2 VSV-G and 10 µg of ΔR8.74 plasmids. HIV-1 viruses were produced by cotransfection with calcium phosphate with HIV-1 LAI (BRU) ΔEnv Virus (NIH) or with the modified versions HIV-1ΔEnvIN_HA_ (41) (kindly provided by Fabrizio Mammano) or HIV-1ΔEnvIN_HA_ΔNef ANCH3 and VSV-G envelope expression plasmid pHCMV-G (VSV-G). The viruses collected from 293T cells 48 h post transfection were ultracentrifuged at 4 °C for 1h at 22,000 rpm. Virus normalizations were performed by p24 ELISA according to the manufacturer’s instructions (Perkin Elmer) and by real time PCR. The titer of each virus has been calculated by qPCR as transducing unit (TU)/mL and then used to calculate the MOI (TU / number of cells). Infectivity has been tested by Beta-galactosidase assay (Merck) activity measured 48 h post infection according to manufacturer’s instructions, using a microplate fluorimeter (Victor, Perkin Elmer). Protein quantification by Bio-Rad protein assay was carried out on the same lysates to normalize the B-gal data for protein content.

## Supporting information

Uninfected HeLa P4R5 cells (control) stably expressing OR-GFP imaged for 71h at the biostation. Cells were imaged every 10 minutes.

## Acknowledgements

We wish to thank Philippe Souque, François Anna and Olivier Gourgette for experimental help, Fabrizio Mammano, Nicoletta Casartelli and Franck Gallardo for scientific discussion and critical reads of the manuscript. We thank Daniela Bruni for help editing the manuscript. We thank Sylvie van der Werf and the department of Virology IP to support the end of the contract of G.B.-R. We thank Fabrizio Mammano, Franck Kirchhoff and NIH reagents program for sharing reagents. Julie Ravel for the inspiration of the study. We thank the members of the Imagopole Platform at the IP. We thank the NIH AIDS Reagents program to support us with precious reagents. This work was funded by the ANRS (Agence Nationale de Recherche sur le SIDA) grant ECTZ4469, the Sidaction/FRM grant n.11291, the Pasteur Institute.

## Author contributions

F.D.N. conceived the study. G.B.-R. and F.D.N. designed the experiments and wrote the manuscript. B.M. and F.D.N. performed live imaging experiments. F.D.N. conceived the HIV-1 ANCHOR system. G.B-R., A.G., F.D.N. and J.K.-L. designed electron microscopy experiments, G.B-R. and A.G. conducted electron microscopy experiments and analysed data. G.B-R. set up and analysed CLEM using immunogold labelling and HIV-1 ANCHOR system. G.B.-R. and F.D.N. performed IF experiments. V.S. performed qPCR experiments and S.F. conducted western blotting, molecular cloning and viral productions. G.B.-R., V.S. and F.D.N. analysed imaging data. F.M., J.K.-L., O.S. and P.C. contributed to stimulating discussions.

