## Supplementary material for "Remodeling of the core leads HIV-1 pre-integration complex in the nucleus of human lymphocytes": movie legends

**Movie S1A.** HeLa P4R5 cells stably expressing OR-GFP infected with HIV-1ΔEnvINHAΔNef ANCH3/VSVG and imaged at the biostation for 71h p.i.. Cells were imaged every 10 minutes.

**Movie S1B.** Uninfected HeLa P4R5 cells (control) stably expressing OR-GFP imaged for 71h at the biostation. Cells were imaged every 10 minutes.

**Movie S2A.** 3D reconstruction by Imaris software of HeLa P4R5 cells stably expressing OR-GFP infected with HIV-1ΔEnvINHAΔNef ANCH3/VSV-G and imaged by spinning disk. Cells were imaged every 20 seconds.

**Movie S2B.** 3D live imaging of HeLa P4R5 cells stably expressing OR-GFP infected with HIV-1ΔEnvINHAΔNef ANCH3/VSV-G 7h post-infection.

**Movie S2C.** 2D live imaging of HeLa P4R5 cells stably expressing OR-GFP infected with HIV-1ΔEnvINHAΔNef ANCH3/VSV-G from 4h post-infection. Cells were imaged every 10 minutes.

**Movie S3A.** HeLa P4R5 cells stably expressing OR-GFP infected with HIV-1ΔEnvINHAΔNef ANCH3/VSVG and imaged at the biostation for 26h p.i. in absence of PF74. Cells were imaged every 5 minutes.

**Movie S3B.** HeLa P4R5 cells stably expressing OR-GFP infected with HIV-1ΔEnvINHAΔNef ANCH3/VSVG and imaged at the biostation for 26h p.i. in presence of low dose of PF74. Cells were imaged every 5 minutes.

**Movie S3C.** HeLa P4R5 cells stably expressing OR-GFP infected with HIV-1ΔEnvINHAΔNef ANCH3/VSVG and imaged at the biostation for 26h p.i. in presence of high dose of PF74 drug. Cells were imaged every 5 minutes.

**Movie S4A.** HeLa P4R5 cells stably expressing OR-GFP infected with HIV-1ΔEnvINHAΔNef ANCH3/VSVG and 3D imaged by confocal at 24h post infection.

**Movie S4B.** HeLa P4R5 cells stably expressing OR-GFP infected with HIV-1ΔEnvINHAΔNef ANCH3/VSVG in presence of 10μM of nevirapine and 3D imaged by confocal at 24h post infection.
